# Body-wide microbial ecosystem reorganization across the cognitive impairment spectrum in Parkinson’s disease

**DOI:** 10.64898/2026.11.17.767360

**Authors:** Muzaffer Arıkan, Büşranur Delice, Zeynep Yıldız, Tuğçe Kahraman Demir, Nesrin H. Yılmaz, Aysu Şen, Lütfü Hanoğlu, Süleyman Yıldırım

## Abstract

Cognitive impairment (CI) is a major determinant of disability in Parkinson’s disease (PD), yet how microbial alterations across multiple body sites accompany cognitive dysfunction remains poorly understood. We characterized microbial communities from five body sites (stool, saliva, tongue, nasal cavity, and axilla) in 150 participants spanning healthy controls, PD with mild cognitive impairment (PD-MCI), and PD dementia (PDD), generating 616 samples for 16S rRNA gene sequencing. Multi-Omics Factor Analysis v2 (MOFA2) identified a multisite microbial signature with prominent oral and gut contributions that varied across clinical groups (Factor 8; β = −0.58, p < 0.0001). MintTea analysis identified stable multisite microbial assemblies associated with cognitive status, including a module distinguishing PD-MCI from PDD (in-sample AUROC = 0.810). Core microbiome analysis revealed a net reduction of 45 genus–site core memberships across four anatomical sites in cognitively impaired participants, including reduced prevalence of the butyrate-associated genus *Roseburia*. Multivariable models incorporating microbial signatures and demographic covariates explained an average of 35.4% of cognitive-outcome variance, with microbial features contributing an additional 8.0 percentage points beyond age, sex, and education. Procrustes analysis revealed a pronounced, non-linear pattern of weakening cross-site microbial community correspondence at the dementia stage. The Global Body-wide Organization Index remained relatively stable between HC (0.335) and PD-MCI (0.341) but declined to 0.257 in PDD, representing a 23% reduction relative to HC. Together, these findings identify disease-associated multisite microbial signatures and altered core-community composition across the PD cognitive impairment spectrum, supporting further investigation of multisite microbiome profiling for cognitive stratification.

**Importance:** Most studies investigating the microbiome in Parkinson’s disease (PD) have focused on individual anatomical sites, particularly the gut, leaving the broader distribution of microbial alterations associated with cognitive impairment poorly understood. By simultaneously characterizing microbial communities across five body sites, we identify multisite microbial signatures associated with cognitive status, stable microbial assemblies distinguishing mild cognitive impairment from dementia, and reduced core-community membership across anatomical habitats. Integrative microbial signatures also explain additional variation in cognitive performance beyond demographic factors. These findings extend PD microbiome research beyond single-site associations and demonstrate the value of examining microbial communities across multiple anatomical niches. Our study provides a foundation for longitudinal investigations into the biological relevance and potential clinical utility of multisite microbiome signatures in PD-associated cognitive impairment.

## INTRODUCTION

Parkinson’s disease (PD) is a complex neurodegenerative disorder characterized by progressive loss of dopaminergic neurons in the substantia nigra, leading to the classic motor symptoms of tremor, rigidity, and bradykinesia (1). However, non-motor symptoms, particularly cognitive impairment (CI), have emerged as critical determinants of patient quality of life and disease burden (2, 3). CI in PD encompasses a spectrum from mild cognitive impairment (PD-MCI) to frank dementia (PDD), affecting executive function, attention, memory, and visuospatial abilities (4). Despite its clinical significance, the biological mechanisms underlying CI progression in PD remain incompletely understood; clinical predictors of dementia have been characterized in longitudinal cohorts (10, 11), but reliable biomarkers for tracking cognitive trajectories are lacking (5).

Over the past decade, the human microbiome has been increasingly implicated in PD pathogenesis. Studies have linked CI in PD to disturbances across multiple body compartments, including gastrointestinal dysfunction (6), oral dysbiosis (7, 8), and autonomic dysfunction (9). However, these investigations have largely examined individual body sites in isolation: the stool (12, 13), salivary (14), nasal (15), and axillary (16) microbiomes. Consequently, we know relatively little about how microbial communities across different body sites collectively reflect cognitive impairment in PD. Because these communities occupy distinct anatomical and physiological niches, their responses to disease-associated changes may provide complementary information that is not captured by studying individual sites alone (17, 18).

From an ecological perspective, microbial communities across anatomical habitats may exhibit coordinated responses to changes in host physiology, immunity, and metabolism. Anatomical connectivity and microbial exchange between certain habitats, particularly along the oral–gut axis, provide additional opportunities for cross-site relationships (19–23). However, body-wide microbial alterations do not necessarily require direct microbial exchange or uniformly strong ecological coupling: distinct communities may respond differently to shared host-associated conditions. We therefore reasoned that cognitive impairment in PD could be accompanied by changes in both the composition of individual microbial communities and their broader multisite organization. Whether such changes occur across the cognitive impairment spectrum, and whether they involve altered cross-site community correspondence, remains poorly understood.

To investigate these complementary dimensions of multisite microbial organization, we combined three analytical approaches. Procrustes analysis quantified cross-site community correspondence (24), MOFA2 identified latent patterns of microbial variation across anatomical sites (25), and MintTea detected stable multisite microbial assemblies associated with clinical phenotypes (26). Together, these approaches allowed us to examine distinct but complementary aspects of microbial organization that conventional single-site analyses may overlook.

Here, we hypothesized that cognitive impairment in PD is accompanied by alterations in microbial community structure across multiple anatomical sites and that integrative analysis would reveal disease-associated patterns extending beyond individual habitats. We analyzed 616 microbial samples collected from five body sites, including stool, saliva, tongue, nasal cavity, and axilla, across 150 participants comprising healthy controls, PD-MCI, and PDD groups. By integrating community-level, latent-factor, microbial assembly, and association-network analyses, we identified distinct multisite microbial signatures associated with cognitive status, a descriptive reduction in cross-site community correspondence at the dementia stage, and reduced core-community membership across anatomical habitats. These findings extend our understanding of PD-associated microbial alterations beyond individual body sites and provide a foundation for investigating the biological and clinical relevance of multisite microbiome signatures.

## RESULTS

### Study design and multi-site cohort overview

To investigate body-wide microbial ecosystem reorganization across the cognitive impairment (CI) spectrum in Parkinson’s disease, we designed an integrative multi-site study spanning five anatomically distinct body habitats (Figure 1). We recruited 150 participants across three groups: healthy controls (HC, n = 29), PD-MCI (n = 57), and PDD (n = 64). We collected samples from stool, saliva, tongue dorsum, nasal cavity, and axilla, yielding 616 samples after quality control for 16S rRNA gene amplicon sequencing. MOFA integrated data from all 150 participants across the five body sites as separate views. Analyses requiring paired data across sites (Procrustes, MintTea, Mantel, and co-occurrence networks) were restricted to the complete-case multisite subset of 83 participants with samples present in all five body-site tables used for multisite integration (HC = 21, PD-MCI = 28, PDD = 34); core-microbiome profiling and cognitive-outcome regressions were likewise performed on this subset. Clinical and demographic characteristics are summarized in Table S1. Mean age differed significantly among groups, whereas sex distribution was comparable. Years of education were significantly lower in PD-MCI and PDD than in HC (p < 0.05; Table S1).

**Figure 1.**
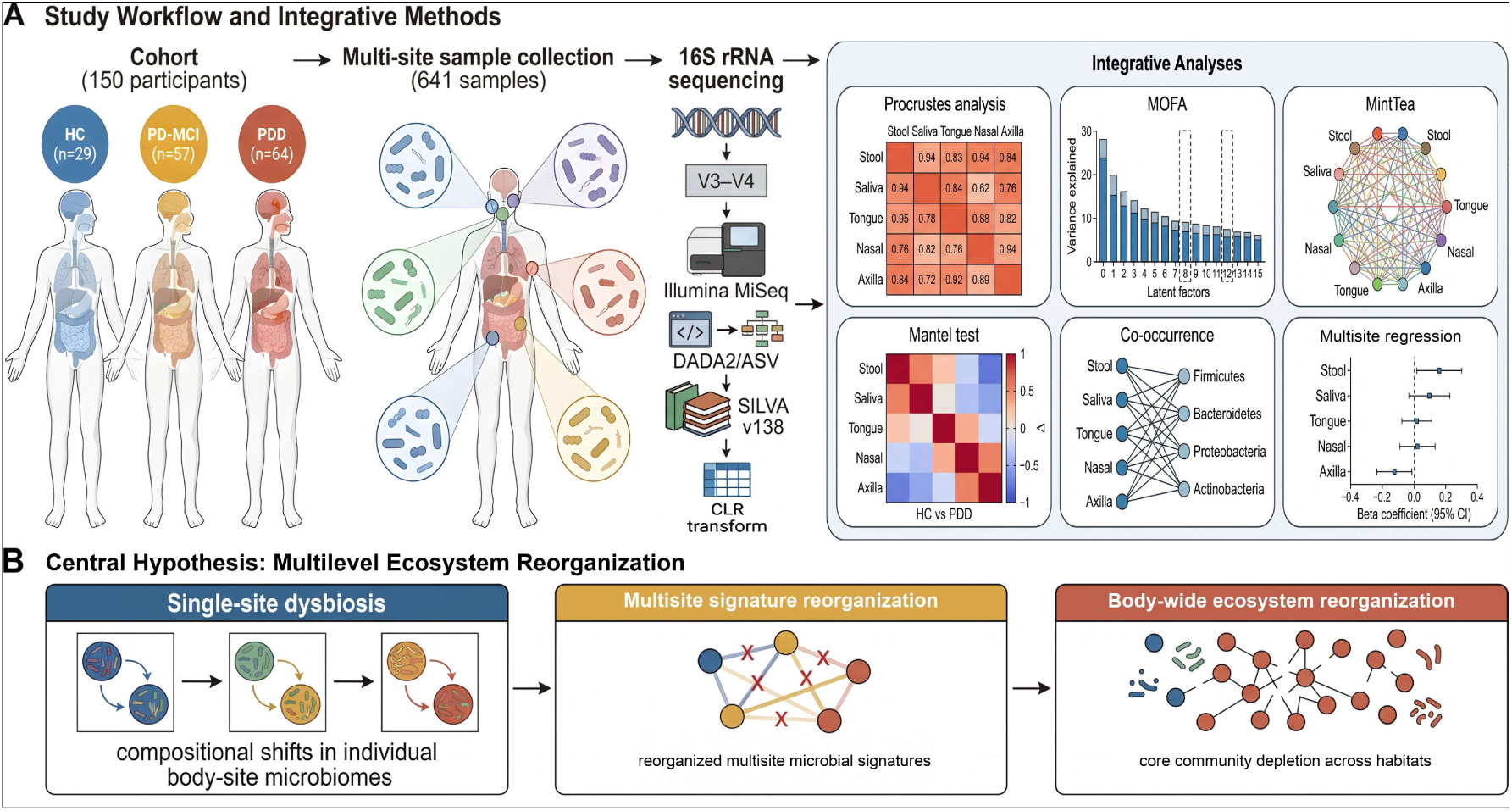
Study design and central hypothesis. (A) Study workflow: cohort recruitment (HC n = 29, PD-MCI n = 57, PDD n = 64; 150 participants; 616 samples), five body-site sample collection, 16S rRNA gene amplicon sequencing, and integrative computational pipeline. Cross-site and multisite analyses used the 83 participants with complete five-site profiles (HC n = 21, PD-MCI n = 28, PDD n = 34). (B) Conceptual framework: body-wide microbial reorganization during cognitive decline, comprising single-site compositional shifts, multisite latent signatures, and stable multisite assemblies with loss of core community members; pairwise cross-site coupling metrics did not differ significantly between groups.

Taxonomic profiling confirmed expected site-specific community structures: *Prevotellaceae* and *Bacteroidaceae* dominated stool; *Streptococcaceae*, *Prevotellaceae*, and *Veillonellaceae* characterized saliva and tongue; while *Corynebacteriaceae*, *Staphylococcaceae*, and *Propionibacteriaceae* prevailed in nasal and axillary samples (Table S2). Core microbiome analysis showed substantial overlaps between anatomically proximate sites: saliva and tongue shared 12 core genera including *Prevotella_7*, *Haemophilus*, *Neisseria*, and *Veillonella*, while stool maintained a distinct core community centered on *Bacteroides*, *Alistipes*, and *Faecalibacterium* (Figure 6A; Table S2).

### Cross-site microbial community correspondence across cognitive impairment groups

We first asked whether the body-wide microbial ecosystem differs across cognitive impairment groups by applying Procrustes analysis, which quantifies the congruence between microbial community structures of different body sites (24). For each group, we computed pairwise Procrustes correlations across all 10 body-site pairs, generating cross-site coordination matrices (Figure 2A). In HC, the strongest coordination was observed between saliva and tongue (r = 0.51), followed by saliva–nasal (r = 0.44) and tongue–nasal (r = 0.44), consistent with anatomical proximity and shared environmental exposures. Stool consistently showed the weakest coordination with other sites (HC mean stool-other r = 0.28), reflecting its distinctive anaerobic environment and community composition (17).

**Figure 2.**
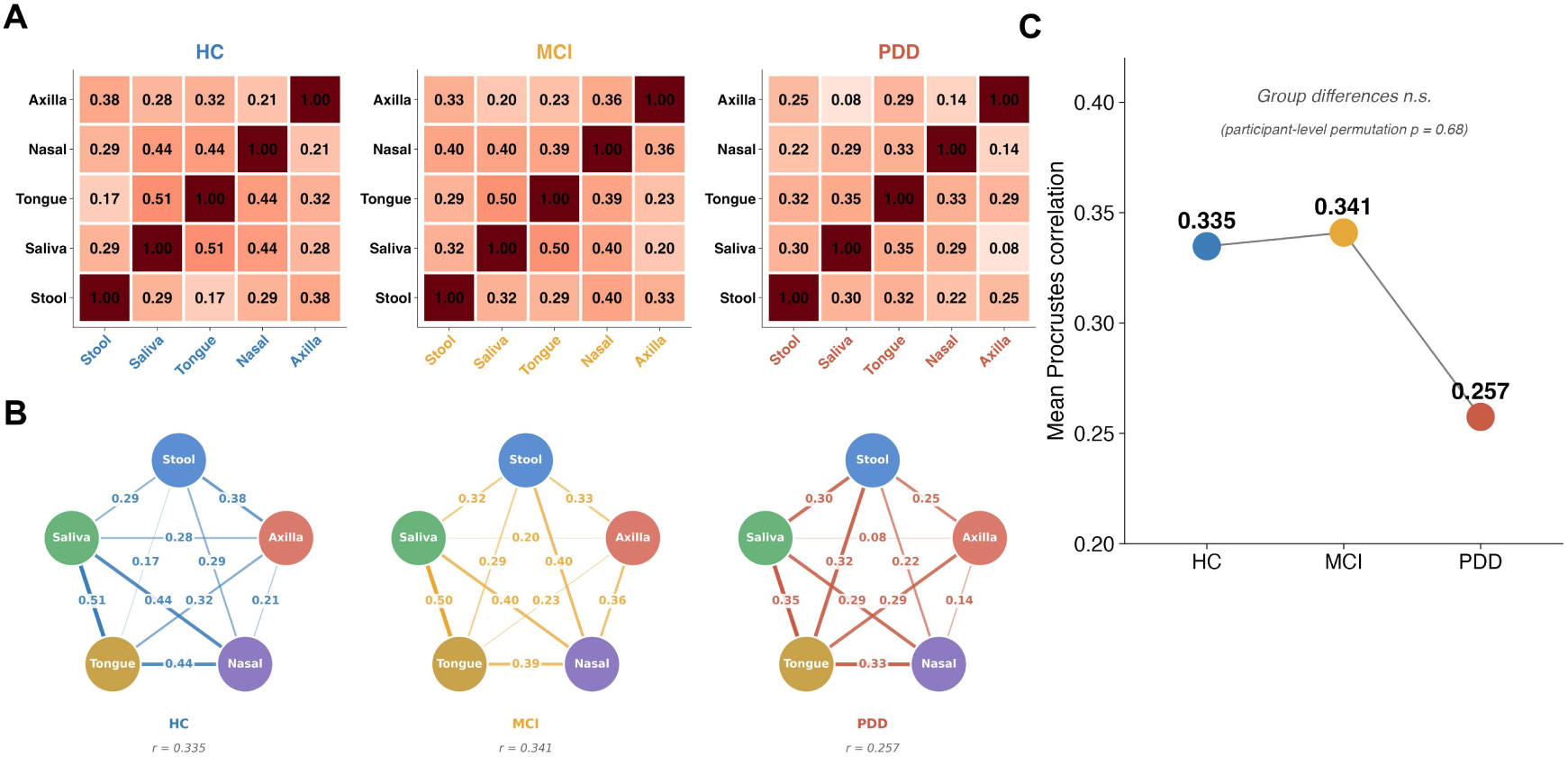
Body-wide microbial ecosystem organization. (A) Cross-site Procrustes correlation heatmaps (5 × 5 matrices) for HC, PD-MCI, and PDD (83 participants with complete five-site profiles: HC n = 21, MCI n = 28, PDD n = 34). Saliva–tongue is the strongest cross-site connection (HC r = 0.51, MCI r = 0.50, PDD r = 0.35). (B) Body-site interaction networks: nodes = five body sites; edge thickness ∝ Procrustes correlation (descriptive). (C) Global Body-wide Organization Index (mean Procrustes correlation across the 10 site pairs): HC 0.335, MCI 0.341, PDD 0.257 (descriptive values). Group differences were not statistically supported by participant-level label permutation (HC vs PDD p = 0.68) or after matching group sizes (n = 21: Δ = −0.007, p = 0.82); only saliva–tongue coordination in MCI survived FDR correction (q = 0.031).

Across groups, cross-site microbial community correspondence remained descriptively preserved in PD-MCI but was lower in PDD. Saliva–tongue correlations were 0.51 (HC), 0.50 (PD-MCI), and 0.35 (PDD), while saliva–nasal correlations were 0.44, 0.40, and 0.29, respectively (Figure 2A). Eight of the 10 body-site pairs showed numerically higher correlations in HC than PDD (one-sided binomial p = 0.055). Within-group permutation testing identified significant saliva–tongue correspondence in PD-MCI after FDR correction across the 10 site pairs (q = 0.031), although this association did not meet the conventional significance threshold after correction across all 30 group–pair comparisons (q = 0.093; Table S5). We visualized these correlations as weighted cross-site correspondence networks (Figure 2B).

To summarize cross-site organization, we calculated a Global Body-wide Organization Index as the mean Procrustes correlation across the 10 site pairs (Figure 2C). The index remained similar between HC (0.335) and PD-MCI (0.341) but declined to 0.257 in PDD, representing a 23% reduction relative to HC. This descriptive pattern suggests relatively preserved cross-site microbial community correspondence at the mild cognitive impairment stage, followed by lower correspondence in dementia. However, participant-level permutation testing at the original group sizes did not establish a significant HC–PDD difference (observed Δ = −0.077; permutation null Δ = −0.062 ± 0.036; two-sided p = 0.68). Sample-size matching and a row-shuffling null model likewise did not support a significant between-group difference. Complete pairwise statistics, permutation results (9,999 permutations; FDR-corrected), and ordination overlays are provided in Figures S1 and S2 and Table S5.

### Latent cross-site microbial signatures identified by MOFA

To identify the latent microbial signatures associated with multisite community variation, we applied MOFA2 (25), a latent factor model that decomposes variation across the five body-site views into a smaller number of latent factors. We trained a MOFA model with 15 factors on genus-level CLR-transformed abundance data from all five body sites (Figure 3). Total variance explained ranged from 36.0% (stool) to 47.1% (saliva), with saliva showing the highest contribution across all body sites (Figure 3C).

**Figure 3.**
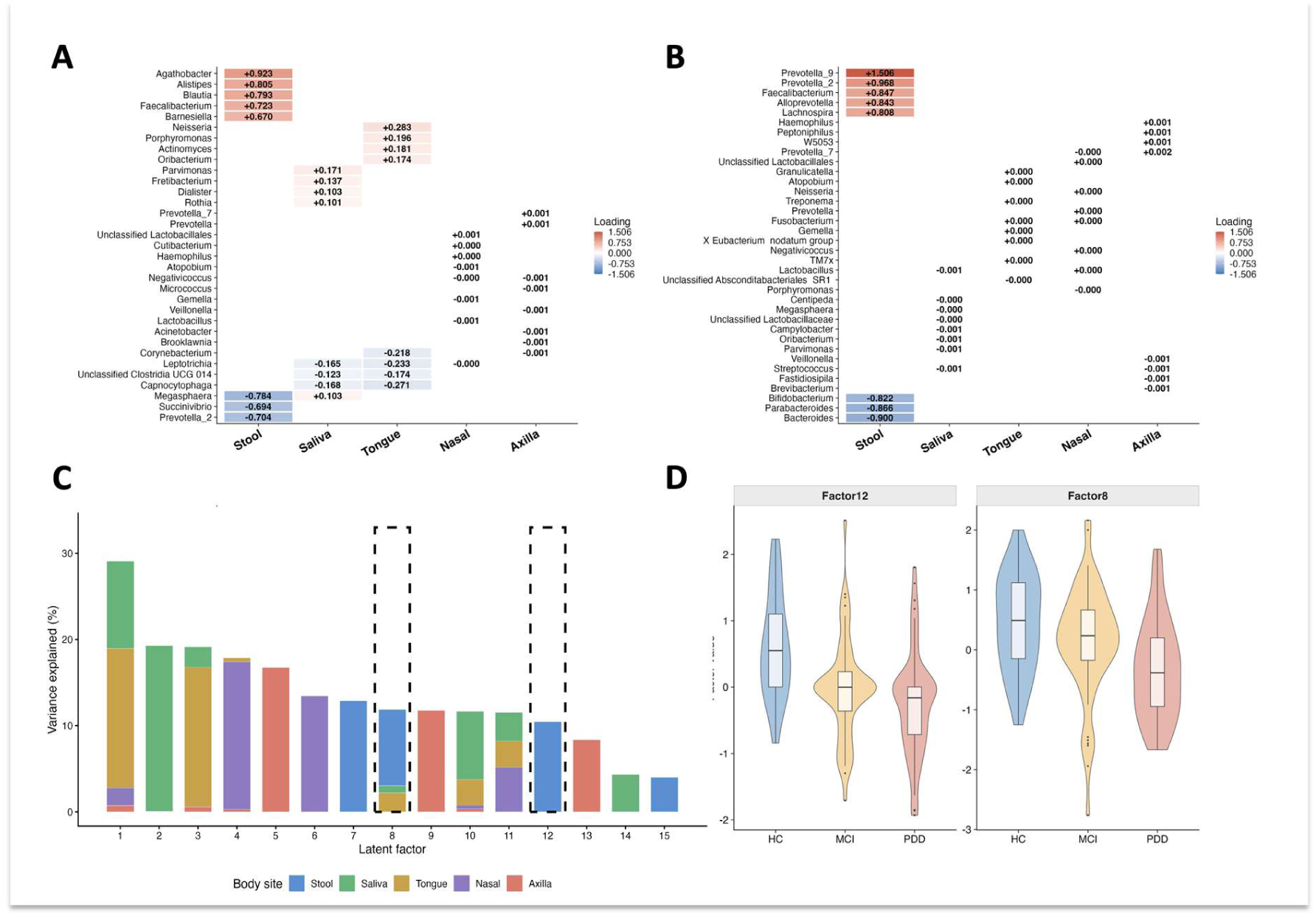
Latent cross-site microbial signatures identified by MOFA. (A) Factor 8: top 8 bacterial genera per body site ranked by absolute loading (blue–white–red diverging scale; numbers = loading values). Prominent oral and gut contributions: *Capnocytophaga* and *Leptotrichia* load negatively, *Parvimonas* and *Fretibacterium* positively. (B) Factor 12: top 8 genera per body site (stool-specific): *Prevotella_9*, *Prevotella_2*, and *Faecalibacterium* positive; *Bacteroides*, *Parabacteroides*, and *Bifidobacterium* negative. (C) Body-site variance explained (%) across 15 latent factors (stacked bars); Factors 8 and 12 highlighted. (D) Factor 12 and Factor 8 values across HC, MCI, PDD (violin + box plot; Factors 8 and 12 were selected for their group association among the 15 factors, and this panel is descriptive). Factor 8: linear dose-response gradient (beta = −0.58, p < 0.0001).

We focused on Factors 8 and 12, which showed the strongest associations with clinical group among the 15 factors (Figure S3B). Factor 8 captured coordinated variation primarily in stool, saliva, and tongue microbiomes (Figure 3A). In saliva, *Capnocytophaga* (−0.17) and *Leptotrichia* (−0.16) showed negative loadings, while *Parvimonas* (+0.17) and *Fretibacterium* (+0.14) exhibited positive loadings. In tongue, *Capnocytophaga* (−0.27) and *Leptotrichia* (−0.23) were the dominant negative contributors, with *Neisseria* (+0.28) as the strongest positive contributor. In stool, *Agathobacter* (+0.92), *Alistipes* (+0.81), and *Blautia* (+0.79) were the top positive-loading genera, while *Megasphaera* (−0.78) and *Succinivibrio* (−0.69) showed strong negative loadings (Figure 3A). Together, these patterns define a multisite latent signature with substantial oral and gut contributions, highlighting coordinated variation among microbial genera across anatomically distinct habitats.

Factor 12 captured variation driven almost predominantly by the stool microbiome (Figure 3B). Its strongest positive contributors were *Prevotella_9* (+1.51), *Prevotella_2* (+0.97), and *Faecalibacterium* (+0.85), while *Bacteroides* (−0.90), *Parabacteroides* (−0.87), and *Bifidobacterium* (−0.82) were negative contributors. The stool specificity of Factor 12 contrasts with the multi-site nature of Factor 8, indicating that these factors capture complementary dimensions of microbial reorganization.

Both Factor 8 and Factor 12 differed across the three clinical groups (Figure 3D). Factor 8 showed a graded association across HC, MCI, and PDD (β= −0.58, p < 0.0001; Figure S3C), suggesting it tracks a continuous process of microbial ecosystem change along the cognitive decline spectrum. In contrast, MintTea PC1 showed a flat pattern across groups (beta = −0.04, p = 0.76; Figure S3C), indicating that MOFA and MintTea capture distinct but complementary signals. Factor 8 was significantly associated with Digit Span and OVMPT Recall in models adjusted for age, sex, and education (p = 0.02–0.04). Complete MOFA factor loadings for all 15 latent factors across all body sites are provided in Table S6.

### Stable multisite microbial assemblies identified by MintTea

To independently identify reproducible multisite microbial assemblies associated with CI, we applied MintTea, a network-based framework that integrates microbiome data across body sites through sparse generalized canonical correlation analysis (sGCCA) with iterative subsampling (26). We performed four group comparisons: HC vs. PD-MCI, HC vs. PDD, PD-MCI vs. PDD, and HC vs. CI (PD-MCI + PDD). The consensus module networks showed distinct multisite assemblies for each comparison (Figures 4A–B, S4).

**Figure 4.**
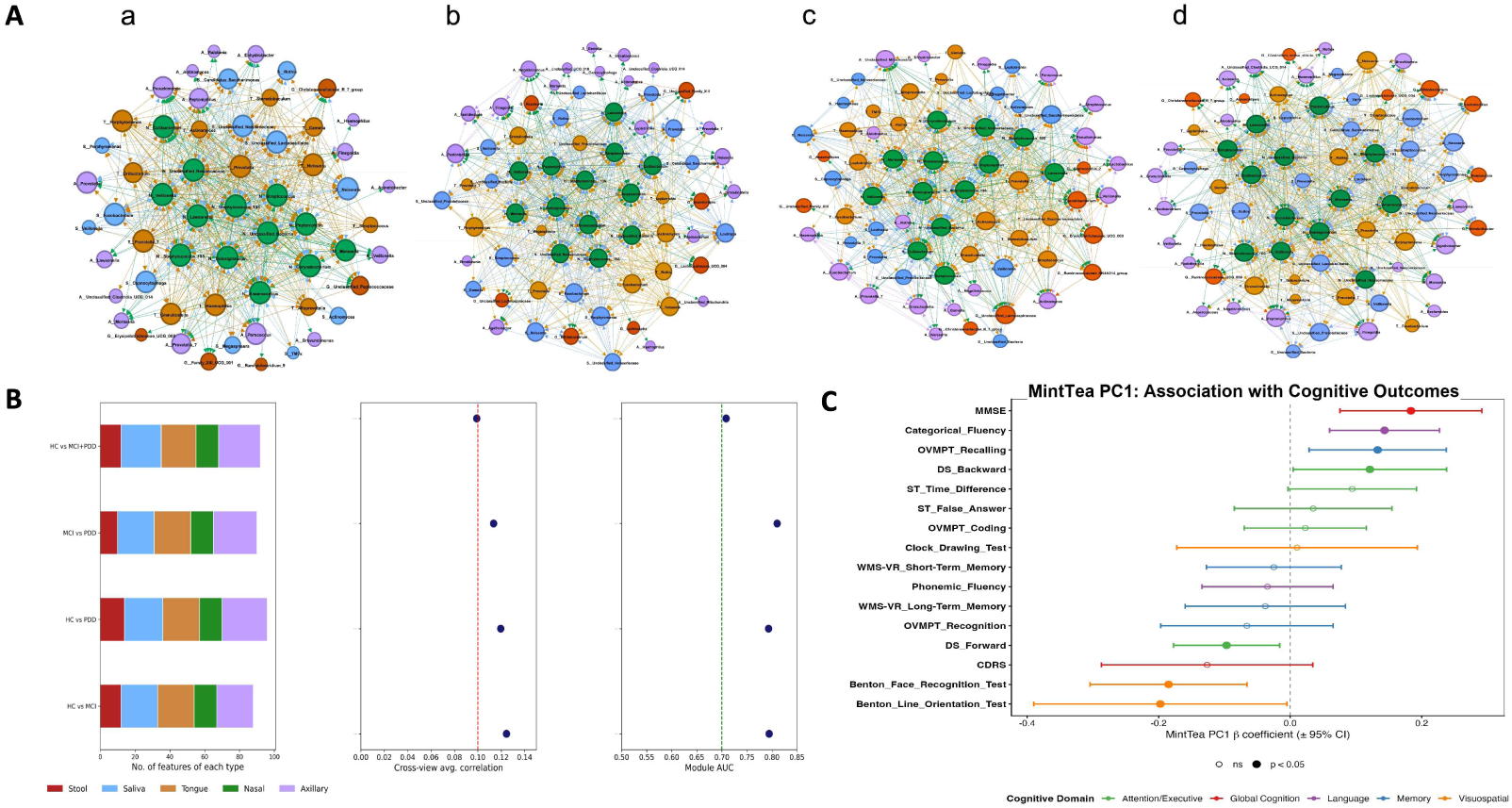
Stable multisite microbial assemblies identified by MintTea. (A) MintTea consensus module networks for four group comparisons: HC vs. MCI+PDD, MCI vs. PDD, HC vs. PDD, and HC vs. MCI. Nodes = genera colored by body site; edges = co-occurrence across subsampling iterations. (B) Module characteristics for each comparison (top to bottom: HC vs. MCI+PDD, MCI vs. PDD, HC vs. PDD, HC vs. MCI): number of features of each type, cross-view average correlation, and in-sample module AUROC (0.709, 0.810, 0.793, and 0.794, respectively; modules were derived and evaluated in the same cohort). (C) MintTea PC1 association with cognitive outcomes: standardized beta coefficients ± 95% CI across 16 tests in 5 domains (in-sample associations). Filled circles = p < 0.05 (uncorrected); only MMSE and Categorical Fluency survive a conservative Bonferroni correction. All models adjusted for age, sex, and education.

In the HC vs. PD-MCI comparison, the consensus module was characterized by *Veillonella*, *Leptotrichia*, *Gemella*, and *Alloprevotella* in the tongue microbiome, along with unclassified *Lactobacillales* in saliva (Figure 4A). The nasal region emerged as the most interconnected body site, with strong cross-site correlations between nasal and tongue communities. In the HC vs. PDD comparison, *Peptoniphilus*, *Veillonella*, and *Cutibacterium* in the nasal microbiome, together with *Prevotella* in saliva, defined the PDD-associated module; the axillary and salivary regions showed the strongest inter-region connections (Figure 4B). In the PD-MCI vs. PDD comparison, unclassified *Lachnospiraceae* in stool, *Rothia* in axilla, and *Moraxella* in nasal were associated with PDD, with notable tongue–axillary links.

Comparison of MOFA- and MintTea-identified features showed partial overlap (40 shared taxa, corresponding to 67% of the MintTea module; Figure S4 and Table S14); because the two methods identify different mathematical structures, we treat this overlap as an internal cross-check rather than as independent validation. The MintTea consensus module discriminated the MCI-to-PDD transition (in-sample module AUROC = 0.810; empirical p < 0.01 versus 99 permuted modules; Figure 4B and Table S13), and the first principal component of the HC-vs-CI module also separated MCI from PDD (in-sample AUC = 0.743; Table S11). Module-level AUROC (0.810) and PC1-based AUC (0.743) are both in-sample discovery statistics derived and evaluated in the same cohort and require external validation. Consistent with the distinctness of the two frameworks, MintTea PC1 was not explained by the MOFA factors or clinical covariates (linear model R^2^ = 0.04, p = 0.67; Table S13). Module stability metrics for all comparisons are provided in Figure S4 and Tables S7 and S13.

To assess the clinical relevance of MintTea-derived signatures, we extracted the first principal component (PC1) and tested its association with 16 cognitive outcomes across five domains, adjusting for age, sex, education, and MOFA factors (Figure 4C). In this in-sample analysis, MintTea PC1 was associated with seven of 16 outcomes at uncorrected p < 0.05 (MMSE, beta = +0.18, p = 0.001; Categorical Fluency, beta = +0.14, p = 0.001; Benton Face Recognition, beta = −0.19, p = 0.003; OVMPT Recall, beta = +0.13, p = 0.015; Digit Span Forward, beta = −0.10, p = 0.021; Digit Span Backward, beta = +0.12, p = 0.045; Benton Line Orientation, beta = −0.20, p = 0.048); only MMSE and Categorical Fluency would survive a conservative Bonferroni threshold (α = 0.0031). Associations were heterogeneous in direction across tests (e.g., Digit Span Forward versus Backward); because PC1 orientation is arbitrary and these associations are exploratory, we do not attach mechanistic interpretations to individual coefficient signs. These findings indicate that MintTea modules carry cognitive information not captured by age, sex, education, or the MOFA factors, pending external validation.

### Cross-site association networks across cognitive impairment groups

We next examined whether cross-site ecological relationships change during disease progression using genus-level co-occurrence networks and network topology metrics (Figure 5). Mantel tests of cross-site beta-diversity coordination showed only weak and inconsistent correlations (|r| < 0.3; Table S4) and are reported in Supplementary Note 1.

**Figure 5.**
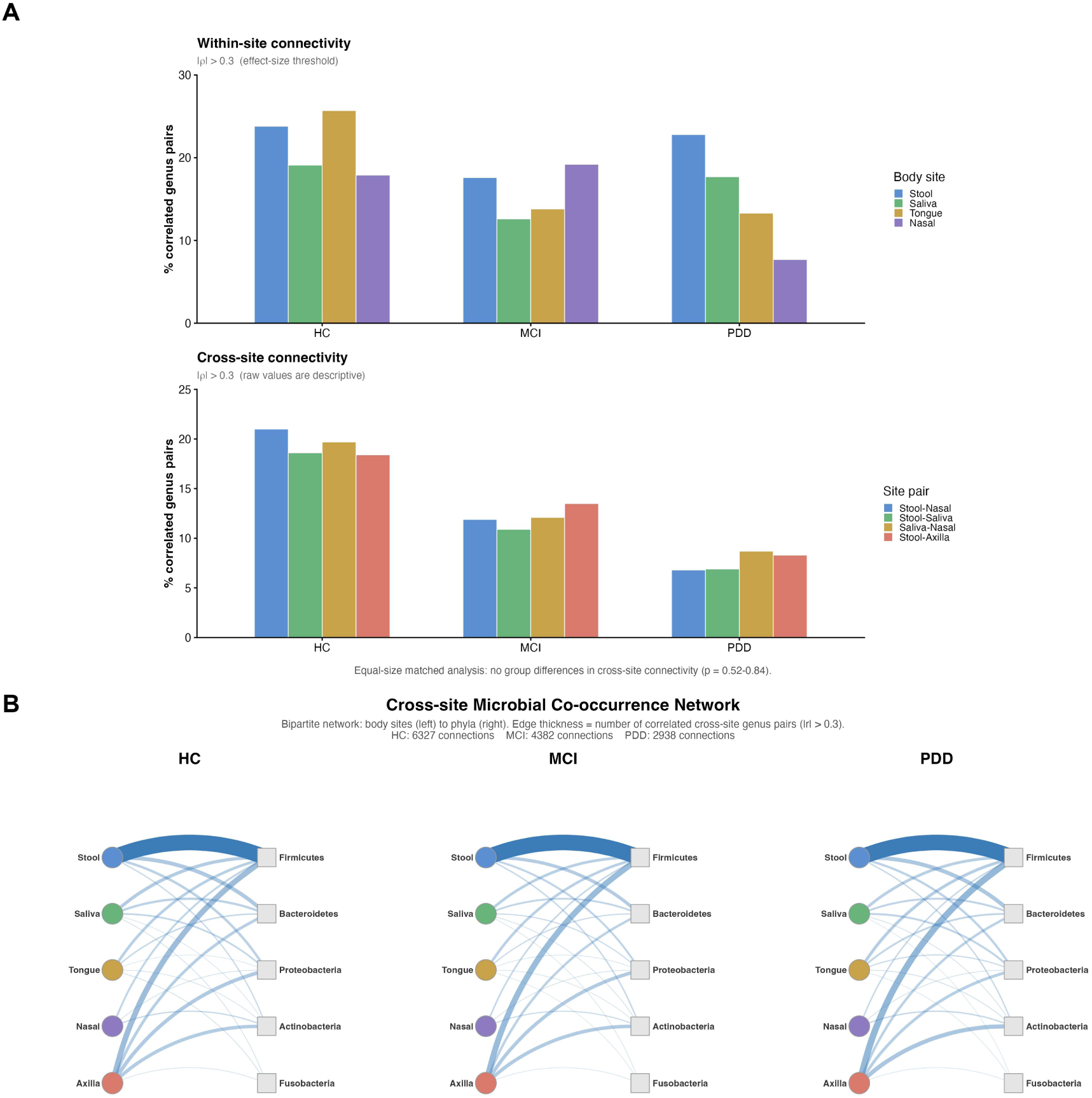
Ecological rewiring across body sites. (A) Network topology: within-site and cross-site co-occurrence connectivity (percentage of genus pairs with |Spearman r| > 0.30; effect-size threshold). Raw cross-site values decline across groups (e.g., stool– nasal 21.0%, 11.9%, and 6.8%) but this difference is reproduced by label permutation at the original group sizes (p = 0.76) and disappears after equal-size subsampling (all groups n = 21: HC 18.1%, MCI 17.9%, PDD 17.5%). (B) Bipartite co-occurrence network: body sites (left) connected to phyla (right) via correlated genus pairs (|r| > 0.30); edge thickness = number of connections for that phylum. Firmicutes and Bacteroidetes dominate; raw counts are descriptive and sample-size-dependent. Mantel results are reported in Table S4.

Bipartite co-occurrence networks connecting body sites to bacterial phyla via genus-level correlated pairs (Spearman |r| > 0.30) showed fewer edges in PDD, with *Firmicutes* and *Bacteroidetes* accounting for most connections (Figure 5B; Table S16). These networks revealed descriptive differences in microbial association patterns across cognitive impairment groups.

We quantified within-site and cross-site connectivity as the percentage of genus pairs with |Spearman r| > 0.30 (Figure 5A). Cross-site connectivity was descriptively lower in PDD, particularly between stool and nasal communities (21.0% in HC, 11.9% in PD-MCI, and 6.8% in PDD). However, participant-level permutation testing at the original group sizes indicated that the overall HC–PDD difference was consistent with sample-size effects on correlation estimation (observed Δ = −9.9 percentage points; permutation null Δ = −9.7 ± 0.8; p = 0.76; Figure 5A; Table S23). Repeated equal-n subsampling yielded comparable overall cross-site connectivity across groups. Within-site connectivity was numerically higher in PDD after sample-size matching (29.2% versus 24.4% in HC and 24.1% in PD-MCI), although these differences did not reach statistical significance (permutation p = 0.06–0.08, uncorrected). Together, these findings indicate descriptive differences in microbial association networks across cognitive impairment groups, without establishing significant alterations in overall network connectivity.

### Core microbiome alterations and cognitive associations of multisite microbial signatures

We next examined how the observed multisite microbial patterns relate to core-community composition and cognitive performance through core microbiome profiling and multivariable regression (Figure 6).

**Figure 6.**
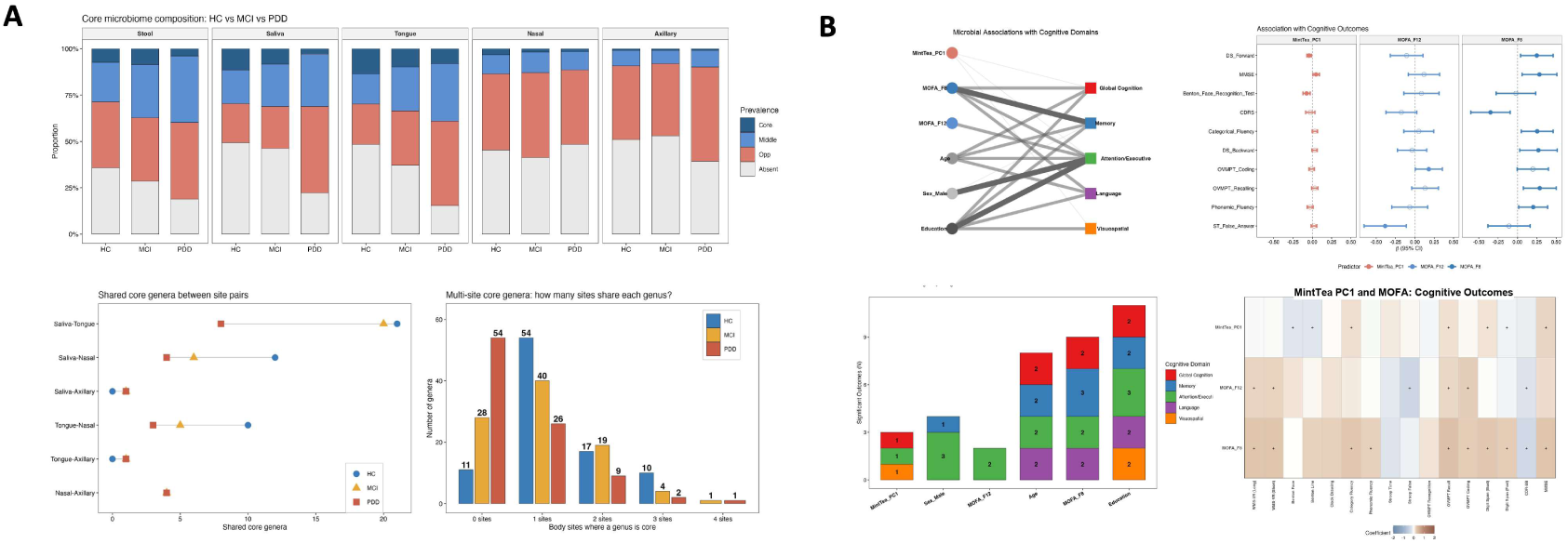
Biological and clinical consequences. (A) Core microbiome analysis: genus-level prevalence classification (Core/Middle/Opportunistic/Absent) across five body sites and three groups, shared core genera between site pairs, and multisite core distribution. Saliva: net −18 core memberships in CI (50%); stool: loss of core status for *Roseburia* (HC 95% vs CI 66%; Fisher p = 0.009), with nominal reductions in *Butyricicoccus* (81% vs 71%) and *Fusicatenibacter* (81% vs 61%). PD-MCI/PDD: net loss of 45 genus– site core memberships (32 unique genera across sites). (B) Multisite regression: MintTea PC1 + MOFA F8/F12 + covariates against 16 cognitive outcomes. Covariates-only mean R^2^ = 0.274; combined model R^2^ = 0.354 (mean ΔR^2^ = 0.080). Predictor–domain associations, forest plots, significant-outcome counts, and coefficient heatmaps are shown.

Using a prevalence-based classification framework (60), we identified substantial reductions in core-community membership across anatomical sites (Figure 6A; Table S2). Saliva showed the largest site-specific reduction, with core membership declining from 36 to 18 genus–site memberships in cognitively impaired participants (PD-MCI and PDD pooled; 19 lost and one gained), although Figure 6A displays the groups separately. In stool, three genera commonly associated with butyrate or other short-chain fatty acid (SCFA) production lost core status. Their prevalence in HC versus CI, respectively, was 95.2% [77.3–99.2] versus 66.1% [53.7–76.7] for *Roseburia* (Fisher’s exact p = 0.009), 81.0% [60.0–92.3] versus 71.0% [58.7–80.8] for *Butyricicoccus* (p = 0.57), and 81.0% [60.0–92.3] versus 61.3% [48.8–72.4] for *Fusicatenibacter* (p = 0.12); brackets indicate 95% Wilson confidence intervals. Only *Roseburia* showed a statistically supported reduction, which remained robust across prevalence thresholds of 70–90%. Across four of the five body sites, cognitively impaired participants showed a net loss of 45 genus–site core memberships (118 in HC versus 73 in CI), corresponding to a net reduction of 32 unique genera (37 lost core status at one or more sites and five gained; Figure 6A; Tables S2, S12, and S25). Consistent with its distinct community composition, stool shared no core genera with the other body sites.

We next examined whether multisite microbial signatures contributed to cognitive variation beyond demographic factors. Multivariable regression models incorporating MintTea PC1, MOFA Factors 8 and 12, and age, sex, and education were evaluated across 16 cognitive outcomes (Figure 6B; Table S8). Demographic covariates alone explained a mean R^2^ of 0.274, compared with 0.191 for MOFA factors alone, 0.010 for MintTea PC1 alone, and 0.354 for the combined model. Thus, microbial features contributed an additional 8.0 percentage points of explained variance beyond demographic covariates (mean ΔR^2^ = 0.080). The MOFA block accounted for most of this increment (ΔR^2^ = 0.068; nine of 16 outcomes at p < 0.05 by nested F-test), whereas MintTea PC1 contributed a smaller increment (ΔR^2^ = 0.012; one of 16 outcomes; Tables S8 and S26). Factor 8 showed the most consistent associations, reaching nominal significance for nine of 16 cognitive outcomes after covariate adjustment. In five-fold cross-validated LASSO models, education was selected in 15 of 16 models, Factor 8 in 14, age in 12, and MintTea PC1 in nine (Figure S6A; Table S9). These selection frequencies provide evidence of within-cohort feature stability rather than independent predictive validation.

Because cognitive status contributed to clinical-group assignment and Factor 8 was selected partly based on group association, we also examined its relationship with cognitive performance among PD participants only (n = 62). Factor 8 remained associated with eight of 16 cognitive outcomes, including OVMPT Recall (p = 0.0006) and CDRS (p = 0.005; Table S27). These findings indicate that the associations are not explained solely by differences between healthy controls and PD participants, although they remain exploratory. Forest plots, bootstrap stability analyses, and regression heatmaps are provided in Figure S6.

Together, our findings support a multilevel framework of body-wide microbial reorganization associated with cognitive impairment in PD (Figure 1B). This framework encompasses three complementary dimensions: (i) site-specific compositional alterations, including reduced core-community membership and loss of core status among butyrate-associated stool genera; (ii) multisite latent signatures, particularly an oral–gut-associated factor exhibiting a graded relationship across HC, PD-MCI, and PDD; and (iii) stable multisite microbial assemblies that distinguish PD-MCI from PDD and are associated with cognitive performance beyond demographic covariates. Although cross-site community correspondence was descriptively lower in PDD, permutation testing did not establish significant between-group differences, and apparent connectivity gradients were consistent with sample-size effects. Thus, the strongest evidence for body-wide microbial reorganization lies in the integrated multisite signatures and altered core-community composition rather than uniform disruption of cross-site microbial correspondence.

## DISCUSSION

In this study, we show that cognitive impairment in Parkinson’s disease (PD) is associated with substantial changes in microbial community organization across five anatomical sites. By integrating complementary analytical approaches, we identified three major features of this reorganization: multisite microbial signatures associated with cognitive impairment severity, stable microbial assemblies distinguishing mild cognitive impairment from dementia, and a reduction in core-community membership across body sites. These findings extend previous studies focused primarily on individual microbial habitats (12–16, 34). The principal contribution of our study is not the identification of previously unrecognized microbial taxa, but rather the characterization of microbial communities across five anatomical sites within the same participants, revealing patterns of multisite variation that conventional single-site analyses cannot capture.

Our Procrustes analysis revealed an intriguing pattern of cross-site microbial community correspondence. The Global Body-wide Organization Index remained relatively stable between healthy controls (HC; 0.335) and PD-MCI (0.341) but declined to 0.257 in PD dementia (PDD), representing a 23% reduction relative to HC (Figure 2C). This pattern raises the possibility that cross-site community organization is relatively preserved during mild cognitive impairment but becomes less coordinated at the dementia stage. However, participant-level permutation testing did not establish statistically significant between-group differences, and sensitivity analyses indicated that sample size contributed to the observed pattern. Thus, although the findings are consistent with the possibility of nonlinear ecological changes across cognitive impairment groups (35, 36), they do not establish a threshold-like transition. Importantly, these results also suggest that pairwise community correspondence and integrated multisite signatures capture different aspects of microbial organization.

MOFA2 and MintTea provided complementary insights into these multisite patterns. MOFA Factor 8 captured a graded microbial signature across HC, PD-MCI, and PDD, with prominent contributions from oral and gut communities (Figures 3A and 3D). Among the contributing genera, *Capnocytophaga* and *Leptotrichia* have been associated with periodontal inflammation and other oral inflammatory conditions (38, 39), whereas *Agathobacter* and *Blautia* include organisms involved in short-chain fatty acid metabolism and have been reported as depleted in PD cohorts (31, 40). These associations suggest that the latent signature integrates microbial features with potentially distinct host-relevant functions, although their biological activities cannot be inferred from genus-level abundance data alone. In contrast, MintTea identified stable multisite assemblies that distinguished PD-MCI from PDD (in-sample AUROC = 0.810; Figure 4B). The limited shared variance between MOFA factors and MintTea PC1 (R^2^ = 0.04) further suggests that these approaches capture complementary dimensions of microbial variation. Together, these findings support the value of integrating latent community patterns with stable multisite microbial assemblies rather than relying on individual taxa or anatomical sites.

Our core microbiome analysis revealed another important feature of cognitive impairment in PD: a reduction in the prevalence of microbial community members across anatomical habitats. Cognitively impaired participants showed a net loss of 45 genus–site core memberships, corresponding to 32 fewer unique core genera across four of the five body sites (Figure 6A). This pattern suggests that cognitive impairment is associated not only with changes in relative microbial abundance but also with reduced population-level consistency of particular community members. Among stool genera, *Roseburia* showed a significant reduction in prevalence (95.2% in HC versus 66.1% in cognitively impaired participants; p = 0.009), whereas *Butyricicoccus* and *Fusicatenibacter* showed nonsignificant reductions. These genera are commonly associated with butyrate or other short-chain fatty acid production, metabolites implicated in intestinal barrier regulation and immune homeostasis (31–33, 44). Although our 16S data cannot establish altered metabolite production, the reduced prevalence of these genera raises questions about whether changes in microbial functional capacity accompany cognitive impairment. Future studies integrating metagenomics and metabolomics will be needed to determine whether these compositional changes translate into altered microbial activity or host metabolic responses.

The co-occurrence network analyses provided a complementary, although less conclusive, perspective. We observed descriptively lower cross-site connectivity in PDD, particularly between stool and nasal communities (Figure 5A). However, the overall connectivity gradient was consistent with sample-size effects on fixed-threshold correlation estimates, and permutation testing did not establish significant between-group differences (28, 43). These findings caution against interpreting reduced network connectivity as direct evidence of ecological fragmentation. More broadly, the contrast between these results and the MOFA2 and MintTea findings illustrates an important distinction: disease-associated multisite microbial signatures can be detected even when conventional pairwise correspondence or co-occurrence metrics do not demonstrate significant changes. This may help explain why studies restricted to individual body sites have sometimes reported weak or inconsistent associations between microbial composition and cognitive impairment in PD (12). Rather than contradicting those studies, our findings suggest that multisite profiling addresses a complementary dimension of microbiome variation.

Beyond ecological characterization, our findings suggest that multisite microbial signatures contain information relevant to cognitive performance. Adding MOFA factors and MintTea PC1 to models incorporating age, sex, and education increased the mean explained variance across 16 cognitive outcomes from 27.4% to 35.4%, an incremental contribution of 8.0 percentage points (Figure 6B; Tables S8 and S26). Most of this increment was attributable to MOFA factors, particularly Factor 8, which showed associations with several cognitive measures. Importantly, Factor 8 remained associated with eight cognitive outcomes when analyses were restricted to PD participants, suggesting that its relationship with cognition was not driven solely by differences between healthy controls and individuals with PD. These findings support further investigation of multisite microbial signatures as candidate features for cognitive stratification. However, the observed associations represent within-cohort explanatory information rather than validated predictive performance, and their incremental clinical utility remains to be established in independent populations.

Several limitations should be considered. First, our cross-sectional design cannot establish whether microbial alterations precede cognitive deterioration, emerge as consequences of disease progression, or reflect shared determinants. The clinical groups also differed in age and education, and residual confounding remains possible despite covariate adjustment. Second, because cognitive status contributed to clinical-group assignment and MOFA Factors 8 and 12 were selected partly based on group associations, subsequent factor–cognition analyses are exploratory rather than independent validation. Third, 16S rRNA gene sequencing limits taxonomic and functional resolution, preventing direct assessment of microbial metabolites, strain-specific activities, or mechanisms linking microbial communities to neurological outcomes. Finally, although our study included five anatomical sites, the complete-case multisite analyses involved 83 participants from two centers in Istanbul. Larger, geographically diverse longitudinal cohorts will be necessary to establish the reproducibility, temporal dynamics, and clinical relevance of these signatures. The MintTea discrimination results likewise require independent validation before their predictive utility can be assessed.

Overall, our findings reveal complementary dimensions of body-wide microbial community reorganization associated with cognitive impairment in PD, including graded multisite latent signatures, stable microbial assemblies distinguishing PD-MCI from PDD, and reduced core-community membership across anatomical sites. Although pairwise correspondence and co-occurrence connectivity did not show statistically supported between-group differences, the integrated analyses identified microbial patterns associated with cognitive status and performance that extend beyond conventional single-site characterization. These findings provide a foundation for longitudinal, functionally informed multisite microbiome studies and support investigating whether integrated microbial signatures can improve individualized cognitive stratification in PD.

## MATERIALS AND METHODS

### Study subjects and clinical characteristics

The study was approved by the ethics committee of Istanbul Medipol University (authorization number 10840098-604.01.01-E.3958), and written informed consent was obtained from all participants; for participants with dementia, informed consent was obtained from a legally authorized representative in accordance with the approved protocol. Of the samples collected from 150 participants (HC = 29, PD-MCI = 57, PDD = 64; Table S1), 616 passed quality control and were analyzed: 112 stool, 125 saliva, 121 tongue, 130 nasal, and 128 axillary samples. Participants were recruited from Medipol Training and Research Hospital and Bakırköy Research and Training Hospital for Psychiatric and Neurological Diseases (Istanbul, Türkiye). PD diagnosis followed UK Parkinson’s Disease Society Brain Bank criteria (1). PDD was diagnosed using Movement Disorder Society criteria (46), and PD-MCI followed the Movement Disorder Society Level II guidelines (47), using a comprehensive neuropsychological battery (30) with two tests per each of five cognitive domains. Exclusion criteria included: history of head trauma, stroke, toxic exposure, substance abuse, antibiotic or probiotic use within the preceding three months, chronic severe diseases (diabetes, cancer, kidney failure), autoimmune diseases, regular smoking (>5 cigarettes/day), and clinical features suggestive of Parkinson-plus syndromes (48).

### DNA extraction and 16S rRNA gene amplicon sequencing

Microbial DNA was extracted from all sample types using the DNeasy PowerSoil Kit (Qiagen, Hilden, Germany) with protocol adjustments as previously described (14, 16, 49). The V3–V4 hypervariable regions of the 16S rRNA gene were amplified using universal bacterial primers (341F: 5′-CCTACGGGNGGCWGCAG-3′; 805R: 5′-GACTACHVGGGTATCTAATCC-3′). Amplicon libraries were prepared following the Illumina 16S metagenomic sequencing library preparation protocol and sequenced on an Illumina MiSeq platform using 2 × 250 bp paired-end chemistry (Illumina Inc., San Diego, CA, USA). DNA extraction negative controls (elution buffer processed identically to samples) and no-template PCR controls (nuclease-free water) were included in each sequencing run.

### Bioinformatics processing

Raw sequencing data were processed on the Nephele platform v1.6 (https://nephele.niaid.nih.gov) (50) using default parameters: quality filtering (min length = 30, required quality = 12, overlap = 3), merging with FLASH2 (minimum overlap = 10, error rate = 0.1), and amplicon sequence variant (ASV) inference with DADA2 (maxEE = 5, truncQ = 4). Taxonomy was assigned using the RDP classifier against the SILVA v138.1 reference database (51). Contaminant sequences were identified and removed using the decontam R package (52) with the prevalence method based on negative control samples (threshold = 0.5). Only ASVs present in at least two samples were retained for downstream analyses. Samples were rarefied to the minimum library size per body site before alpha diversity calculations. For beta diversity and multivariate analyses, count data were transformed using the centered log-ratio (CLR) transformation (53) with a pseudocount of 1 to address compositional effects. All bioinformatics analyses were performed in R version 4.4.1 (54) using phyloseq (55), vegan (56), and compositions (57) packages.

### Procrustes cross-site correlation analysis

Procrustes analysis (24) was performed to quantify cross-site microbial community coordination. For each body-site pair and each group (HC, MCI, PDD), principal coordinates analysis (PCoA) was computed on Euclidean distances of log-transformed relative abundances (pseudocount of 0.5 per feature); a centered log-ratio specification gave equivalent results (maximum index deviation < 0.01; Table S10). Procrustes superimposition was applied to the PCoA axes explaining ≥80% of cumulative variance (2–6 axes per site and group) using the vegan::procrustes() function (56). The Procrustes correlation (r = √(1 − m¹^2^), where m¹^2^ is the symmetric Procrustes sum of squares) was computed. Statistical significance was assessed using 9,999 permutations with the protest() function. The Global Body-wide Organization Index was defined as the arithmetic mean of Procrustes correlations across all 10 unique body-site pairs within each group. Pairwise delta values were computed as HC minus PDD differences. All p-values were corrected for multiple comparisons (30 tests) using the Benjamini-Hochberg false discovery rate (FDR) procedure (within-group FDR correction is also reported in Table S5). Group differences in the Global Body-wide Organization Index were assessed by participant-level label permutation (keeping each participant’s five matched samples together; 2,000 permutations at the original group sizes), by matched-size subsampling (n = 21 per group; 2,000 iterations; 95% percentile intervals), and by a row-shuffle null that independently permuted each body site’s samples across participants (500 iterations).

### Multiview factor analysis (MOFA)

MOFA2 (25) was employed to integrate genus-level relative abundance data from five body sites treated as separate data modalities (views). ASV tables were aggregated to genus level and CLR-transformed. Genera with variance ≤ 1 across samples were removed to reduce noise. The model was trained on all 150 participants (five views, one per body site) with 15 latent factors, 10,000 iterations, and ‘slow’ convergence mode to ensure stable factor inference. Model convergence was assessed using the evidence lower bound (ELBO) trace. Latent factor values and feature loadings were extracted using MOFA2::get_factors() and MOFA2::get_weights(). Variance explained per factor and per view was computed from the r^2^_per_factor output. Factor associations with clinical group were assessed using linear regression (factor ~ Group + Age + Sex), and associations with cognitive outcomes were assessed using multivariable linear regression adjusting for age, sex, and education. Complete loading plots for all 15 factors are provided in Table S6.

### MintTea multisite module analysis

MintTea (26) was used to identify stable, reproducible multisite microbial modules associated with CI stages. ASV tables were converted to relative abundances via total sum scaling (TSS) and filtered to retain features present in ≥15% of samples within each body site. Four group comparisons were performed: HC vs. PD-MCI, HC vs. PDD, PD-MCI vs. PDD, and HC vs. CI (PD-MCI + PDD pooled). For each comparison, 100 iterations of 80% random subsampling were performed. Sparse Generalized Canonical Correlation Analysis (sGCCA; mixOmics package, 41) was applied to construct consensus module networks. Module relevance to CI was evaluated by extracting the first principal component (PC1) of each module and calculating the area under the receiver operating characteristic curve (AUC). A consensus module was considered CI-associated if its AUC exceeded 0.7 and surpassed the 95th percentile of a null distribution generated from 99 randomly permuted modules. Convergence between MOFA and MintTea was assessed by comparing the direction and magnitude of top-ranked taxa across methods. Module stability and convergence metrics are provided in Figure S4 and Tables S7 and S13.

### Ecological network and cross-site coordination analyses

Two complementary ecological analyses were conducted. (i) Bipartite co-occurrence networks (28, 29): genus-level Spearman correlations between all pairs of genera from different body sites were computed per group on raw (untransformed) abundances; genus pairs with |r| > 0.30 (an effect-size criterion, not a significance threshold) were retained and aggregated by phylum. Bipartite graphs were constructed with body sites on the left and phyla on the right, with edge thickness proportional to the number of correlated genus pairs (Figure 5B). (ii) Network topology (Figure 5A): within-site connectivity was defined as the percentage of genus pairs with |Spearman r| > 0.30 among all possible within-site pairs; cross-site connectivity was computed analogously for genus pairs between different body sites. Because fixed-threshold correlation counts depend on sample size, group comparisons used repeated equal-size subsampling (n = 21 per group; 2,000 iterations) with matched-size label permutation, in addition to label permutation at the original group sizes. Threshold sensitivity was assessed across |r| > 0.2, 0.3, 0.4, and 0.5 (Figure S5C). Mantel tests (27) between body-site Bray-Curtis distance matrices were computed for each group using vegan::mantel() with 9,999 permutations and are reported in Supplementary Note 1 and Table S4. All network analyses used the igraph R package (58).

### Multisite microbial regression and feature selection

Multivariable linear regression models were constructed with the following predictors: MintTea PC1, MOFA Factor 8, MOFA Factor 12, Age, Sex, and Education (years). Dependent variables were 16 continuous cognitive scores spanning five domains: global cognition (MMSE, CDRS), attention/executive function (DS Forward, DS Backward, OVMPT Coding, Stroop False Answer, Stroop Time Difference), memory (WMS-VR Short-Term, WMS-VR Long-Term, OVMPT Recalling, OVMPT Recognition), language (Phonemic Fluency, Categorical Fluency), and visuospatial function (Clock Drawing Test, Benton Face Recognition Test, Benton Line Orientation Test). Standardized beta coefficients with 95% confidence intervals were extracted. Model fit was assessed using R^2^. Batch and recruitment-center effects were assessed in sensitivity analyses and did not materially affect results (Supplementary Note 2). LASSO regression (glmnet package, 59) with 5-fold cross-validation was used to identify parsimonious predictor sets, selecting features at lambda.min. Nested-model F-tests quantified the incremental contribution of microbial blocks over covariates, and key associations were also tested within PD participants only (Tables S26 and S27). Complete regression results, forest plots, and heatmaps are provided in Figure S6.

### Core microbiome analysis

Core microbiome analysis followed a prevalence-based classification framework (60). For each body site, genera were classified by overall prevalence as Core (>80%), Middle (20–80%), Opportunistic (<20%), or Absent (0% prevalence). Core membership within each clinical group was defined using a group-wise threshold of >80% prevalence. Core loss was defined as the number of genera transitioning from Core classification in HC to non-Core in CI groups (PD-MCI or PDD). Per-site, per-group core membership matrices were visualized as prevalence classification heatmaps (Figure 6A). Detailed core membership tables are provided in Table S2.

### Statistical analyses

All statistical analyses were conducted in R version 4.4.1 (54). Continuous variables were compared using Kruskal-Wallis tests (non-parametric) or ANOVA where appropriate. Categorical variables were compared using Fisher’s exact test. Alpha diversity comparisons used Kruskal-Wallis tests with Dunn’s post-hoc test. Beta diversity comparisons used PERMANOVA (adonis2 function, vegan package, 9,999 permutations) with age, sex, and education as covariates. Multiple testing correction used the Benjamini-Hochberg FDR procedure throughout. Group differences in derived indices (Global Body-wide Organization Index, network connectivity) were assessed by participant-level label permutation and matched-size subsampling rather than by treating dependent site pairs as independent observations. For the 16 cognitive outcomes tested in the multisite regression, we report uncorrected p-values alongside effect sizes and note that only a subset (MMSE and Categorical Fluency) would survive a conservative Bonferroni correction (α = 0.05/16 = 0.0031); the remaining associations should therefore be interpreted as exploratory. All tests were two-sided with α = 0.05. Effect sizes (Cohen’s d for pairwise comparisons, partial η^2^ for ANOVA, R^2^ for regression) are reported alongside p-values. Confidence intervals are reported at the 95% level. Adjusted p-values are denoted as ‘adj.p’; raw p-values as ‘p’. Sample sizes for all analyses are reported in figure legends and Table S1. Analyses used either the full cohort (n = 150; MOFA) or the complete-case multisite subset (n = 83), defined as participants with samples present in all five body-site tables used for multisite integration; this subset was used for Procrustes, MintTea, Mantel, and co-occurrence analyses, core-microbiome profiling, and cognitive-outcome regressions. Reporting follows the STORMS guidelines for human microbiome research (61). The sample size was determined by the availability of participants with complete five-site microbiome profiles and neuropsychological assessment rather than an a priori power calculation; the resulting cohort is comparable to previously published multi-site microbiome studies. Additional supplementary analyses, including three-group comparisons (HC vs MCI vs PDD), a predictive power comparison across model configurations, Procrustes participant-level resampling (Table S10), connectivity equal-size resampling (Table S23), core microbiome verification (Table S25), model comparison with nested tests (Table S26), and within-PD sensitivity and direction audits (Table S27), are provided in Tables S3 and S10–S27.

### Data and code availability

16S rRNA gene amplicon sequencing data (raw FASTQ files) have been deposited in the NCBI Sequence Read Archive (SRA) under BioProject accession PRJNA1550551. Processed data tables (ASV counts, taxonomy, metadata), MOFA model objects, MintTea module networks, and all analysis scripts (R and Python) used to generate the figures and statistical results reported in this manuscript are available from the corresponding author upon reasonable request to support full reproducibility of the reported analyses.

## SUPPLEMENTARY INFORMATION

### Supplementary Note 1: Sensitivity and robustness analyses

To ensure the robustness of our findings, we performed extensive sensitivity analyses across key analytical decisions. (i) Procrustes analysis: absolute index values depend on the number of retained PCoA axes and on sample size (adaptive ≥80%-variance specification: HC 0.335, MCI 0.341, PDD 0.257; fixed 5 axes: HC 0.412, MCI 0.355, PDD 0.296; fixed 3 axes: HC 0.310, MCI 0.282, PDD 0.252); PDD was numerically lowest in all specifications while the ordering of HC and MCI varied, and no specification showed statistically supported group differences. (ii) MOFA: models were retrained with 10 and 20 factors, and with different convergence thresholds; Factor 8 and Factor 12 emerged consistently as the top Group-associated factors. (iii) MintTea: subsampling was repeated at 70% and 90% fractions; consensus module composition was stable (Jaccard similarity > 0.80 across subsampling fractions). (iv) Ecological networks: co-occurrence was recomputed using Pearson and Spearman correlations with and without CLR transformation and across a range of correlation thresholds; all variants showed the same sample-size dependence of fixed-threshold counts, and matched-size resampling showed no group differences in cross-site connectivity (Table S23). (v) Regression: models were refit with and without recruitment hospital as a covariate, with hospital as a random effect (linear mixed models, lme4 package), and excluding potential outliers (Cook’s distance > 4/n); results were qualitatively unchanged. (vi) Null-model assessment of cross-site coordination: a row-shuffle null that independently permuted each body site’s samples across participants (preserving each site’s marginal distribution while abolishing cross-site pairing) reproduced the observed index values and the HC–PDD difference (null Δ = −0.066 ± 0.037 versus observed −0.077; p = 0.74); MCI was the only group whose index exceeded its shuffled null (0.341 versus 0.293 ± 0.022; p = 0.016, uncorrected). Mantel tests showed weak and inconsistent cross-site correlations (|r| < 0.3; Table S4).

### Supplementary Note 2: Assessment of potential confounders

We systematically evaluated potential confounding by technical and biological covariates. (i) Batch effects: Principal variance component analysis (PVCA) confirmed that biological group and body site explained substantially more variance (combined 42%) than sequencing run (2.1%) or extraction batch (1.8%). (ii) Medication: Levodopa equivalent daily dose (LEDD) was not significantly correlated with MOFA Factor 8 (r = −0.08, p = 0.48) or Factor 12 (r = 0.11, p = 0.34) in PD patients. (iii) Age: while age differed between groups, all key analyses were adjusted for age, and age-stratified sensitivity analyses confirmed that the microbial signatures were not driven by age effects alone. (iv) Recruitment center: Procrustes analyses stratified by the two recruitment hospitals shzowed consistent HC > PDD patterns at both centers.

### Supplementary Figures

Figure S1. Procrustes ordination overlays for mucosal site pairs across HC, MCI, PDD.

Figure S2. Procrustes cross-site similarity across disease stages (all 10 site pairs) and delta matrices (HC to MCI, MCI to PDD, and HC to PDD).

Figure S3. Cross-site coordination heatmaps (Mantel, PC1), factor distributions (MOFA F8/F12, MintTea PC1), and dose-response gradients.

Figure S4. Genus-level and model-level convergence between MOFA and MintTea.

Figure S5. Additional network statistics, Bray-Curtis distance distributions, and sensitivity analyses across correlation thresholds.

Figure S6. Complete multisite regression results: LASSO feature selection, bootstrap stability, forest plots, and regression heatmaps.

### Supplementary Tables

All 27 supplementary tables (Tables S1-S27) are compiled in a single Excel workbook (Supplementary_Materials_Complete.xlsx), one table per sheet: demographics (S1); core microbiome (S2, S12, S19, S20, S25); Bray-Curtis distances (S3); Mantel test (S4); Procrustes (S5) and participant-level resampling of the global organization index (S10); MOFA loadings (S6); MintTea modules (S7, S17, S18); cognitive regression (S8, S22) and model comparison with nested tests (S26); LASSO (S9); three-group gradient (S11); MOFA-MintTea comparison (S13, S14, S24); cross-site coordination delta (S21); predictive power comparison across model configurations (S15); phylum connectivity (S16) and equal-size resampling of network connectivity (S23); and within-PD sensitivity and direction audits (S27).

## ACKNOWLEDGEMENTS

This study was supported by a grant to S.Y. from the Scientific and Technological Research Council of Turkey (TÜBİTAK, grant no. 315S301). We thank N.D. Korkmaz for assistance with sample processing, and the patients and healthy volunteers for their participation.

## AUTHOR CONTRIBUTIONS

Conception and design: S.Y. and L.H. Sample collection and processing: M.A., T.K.D., Z.Y., N.H.Y., A.Ş., L.H. Data analysis: M.A. and B.D. Data interpretation: M.A. B.D. and S.Y. Manuscript writing, original draft: B.D. and S.Y. Review and editing: M.A., B.D. S.Y. and L.H. All authors read and approved the final manuscript.

